# A common framework for the problem of deriving estimates of dynamic functional brain connectivity

**DOI:** 10.1101/215772

**Authors:** William Hedley Thompson, Peter Fransson

## Abstract

The research field of dynamic functional connectivity explores the temporal properties of brain connectivity. To date, many methods have been proposed, which are based on quite different assumptions. In order to understand in which way the results from different techniques can be compared to each other, it is useful to be able to formulate them within a common theoretical framework. In this study, we describe such a framework that is suitable for many of the dynamic functional connectivity methods that have been proposed. Our overall intention was to derive a theoretical framework that was constructed such that a wide variety of dynamic functional connectivity techniques could be expressed and evaluated within the same framework. At the same time, care was given to the fact that key features of each technique could be easily illustrated within the framework and thus highlighting critical assumptions that are made. We aimed to create a common framework which should serve to assist comparisons between different analytical methods for dynamic functional brain connectivity and promote an understanding of their methodological advantages as well as potential drawbacks.

**Highlights:** Different approaches to compute dynamic functional brain connectivity have been proposed, each with their own assumptions.

We present a theoretical framework that encompasses a large majority of proposed methods.

Our common framework facilitates comparisons between different methods and illustrates their underlying assumptions.

## Introduction

Investigations of dynamical functional connectivity (DFC) in the human brain has become achievable owing to recent methodological developments. Foremost, variations on the so called sliding window method have extensively been employed, in which signal co-variance is computed for consecutive segments of the BOLD fMRI signal intensity time series (1–4). Although variations of the sliding window method have gained popularity among the possible routes to estimate DFC, techniques based on other metrics have emerged, such as temporal ICA (5), clustering (6,7), temporal derivatives (8) and change-point detection (9). Advantages as well potential caveats for proposed DFC methods have been outlined and discussed in several reviews and commentaries (2,10,11) as well as their potential to provide a detailed map of the dynamics at a whole-brain level (12–15).

The diversity of proposed strategies for DFC is a sign of methodological development and progress within the field. But the current state of affairs also makes it harder for researchers to be able to understand, interpret and compare results pertaining to dynamic brain connectivity that have been derived by different DFC analytical methods. Currently, there exist no common framework for the theoretical basis of dynamic brain connectivity. This makes it difficult to understand which assumptions that are made either implicitly or explicitly for different DFC methods.

We believe that working towards the goal of a putative common framework for DFC methods would be beneficial for several reasons. First, it would allow researchers to understand which assumptions that are being made for their DFC analyses. Second, it would provide the current manifold of suggested DFC methodologies with a common scaffolding that would serve to assist in validating and comparing different methods. Hence, a common framework would provide a theoretical basis for understanding putative methodologically driven results in dynamic brain connectivity.

The aim of this paper was to find a common ground for many of the proposed DFC analysis strategies. However, we cannot claim that the framework presented here is exhaustive and encompasses all methods, nor do we in this theoretical work present quantitative comparisons and/or validation of previously proposed DFC methods.

## Methods

The data used here was included for the sole purpose of illustration of the concepts described and not intended to provide a basis for a quantitative comparison of the performance of different DFC methods. For this reason, we selected a temporal segment of 50 image volumes of BOLD fMRI signal data extracted from three ROIs in a single subject (Figure 1A). Three regions were selected instead of only two to illustrate the idea that some methods can utilize a state space when handling multiple time series. The data originated from the Beijing eyes open/eyes closed dataset that is publicly available (6) and we have previously reported the preprocessing steps taken (16). We have here chosen to illustrate the difference between different approaches to DFC estimation based on empirical fMRI data rather than on simulated data. This was done with the intent to show that the choice of method to investigate DFC in the brain do have consequences when applied to actual acquired fMRI data and not just in simulated scenarios. All DFC analysis after image preprocessing was done in python 3 using the packages NumPy (17), SciPy (18), Statsmodels (https://github.com/statsmodels/statsmodels], Matplotlib (19), and Teneto (20).

**Figure 1:**
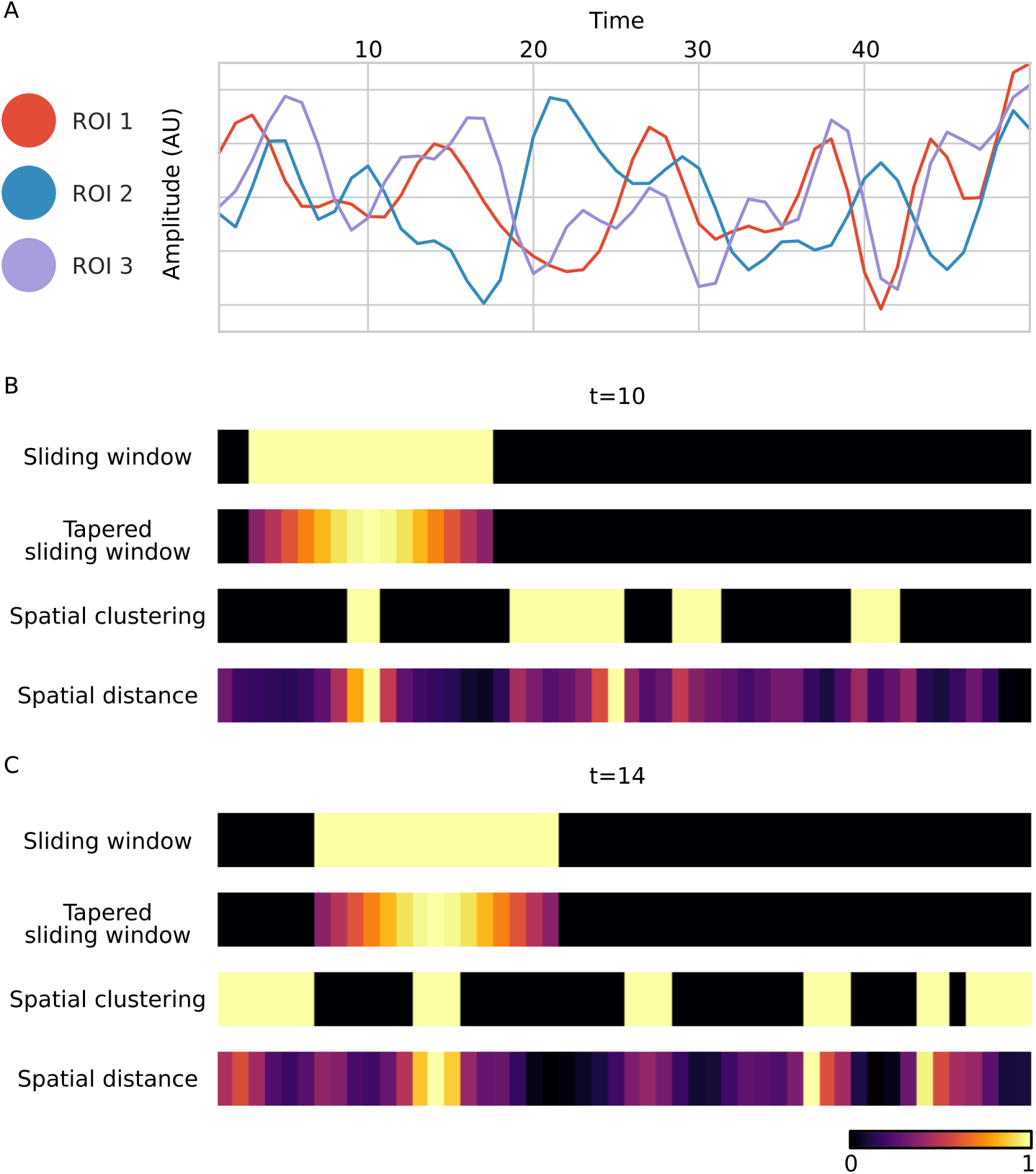
Illustration of the weight-vector formulation for different methods to estimate co-variance in dynamic functional brain connectivity. (A) A sample of three time-series extracted from a single BOLD fMRI resting-state session. (B) Representations of the weight-vectors used for computing DFC for four different methods at *t* = 10. The sliding window size was set to 15 and the tapered sliding window used a Gaussian distribution. The spatial clustering was done by the k-means algorithm (k=3) and for the spatial distance method we used the Euclidean distance. (C) Same as (B) but for *t* = 14.

The static functional connectivity between the three selected BOLD signal time-series were as follows: ROI1 and ROI2: r=−0.099, p=0.495; ROI1 and ROI3: r=0.659, p<0.001; ROI2 and ROI3: r=−0.219, p=0.126. As stated above, the ground truth of DFC for the selected time-series is unknown. However, our main intention in this paper was to show that different assumptions regarding DFC will lead to different estimates of connectivity. A statistical comparison of the performance for different DFC methods was not the primary aim of this paper. However, we have recently performed a detailed simulation and comparison study of current methods to estimate DFC, and we refer the interested reader to (21).

## Results

We observe that for the large majority of DFC methods published so far, a prominent example being the sliding window method, the estimation of co-variability between brain regions employs the Pearson correlation. This fact makes correlation methods a good starting point to formulate a common theoretical framework. To start with, an estimate of the covariance between two variables (i.e. how much they vary together) requires multiple observations. This simple statement also holds the crux of the problem of DFC. Namely, to estimate the connectivity at time-point *t*, we need to take into account more time-points than just *t* itself. This becomes problematic when the ultimate aim is to obtain a unique connectivity estimate at every time-point *t*.

Suppose that we have extracted the signal time-series for brain regions *A* and *B* as *A*_1_, *A*_2_…*A*_*T*_ and *B*_1_, *B*_2_…*B*_*T*_ at [1, 2, 3…*T*]. We are then interested in a subset of time-points, *S*_*t*_, such that *S*_*t*_ ⊂ *T*. The selected subset aims to inform us about the connectivity that happens at *t*. Then, as we will show later, the fundamental assumption and question for all DFC methods is what can be said to be a reasonable assumption in how we choose *S*_*t*_?

As alluded to previously, many DFC methods based on covariance measures use the Pearson correlation coefficient to quantify covariance. This means that the Pearson correlation coefficient at every time-point *t* can be expressed as:

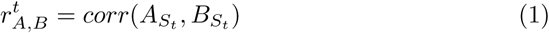

This is simply to say that the degree of correlation between regions *A* and *B* at time-point *t* is the correlation calculated for the subset of time-points included in *S*_*t*_.

Eq. 1 can be illustrated using the sliding window method (without using a taper). In this case, the selection of *S*_*t*_ consists of those time-points adjacent in time with respect to *t*. For example, if the window length is 5, we would have *S*_*t*_ = *t* − 2, *t* − 1, *t*, *t* + 1, *t* + 2 which would then be used to estimate the signal covariance between regions *A* and *B* at time *t*. Thus, the fundamental assumption of the sliding window method is that adjacent time-points close to *t* can be used to estimate connectivity at *t*.

Another way to formulate the information in *S*_*t*_, which contains time-points that are to be included, is to create a weight-vector for all time points *t* (*w*^*t*^) which has a length of *T*. In the sliding window example, the individual weights in the weight-vector will be set 0 for time-point that are not included in *S*_*t*_ and to 1 if they are included in *S*_*t*_ as follows

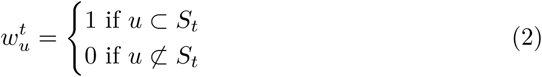

So, the next question is what possible advantage does the introduction of weight-vectors in this context bring? To answer that question, we start by changing the Pearson correlation in eq. 1 to the weighted Pearson correlation with the weights *w*^*t*^ included as a weight-vector as follows

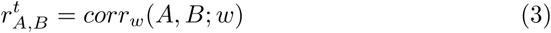

The switch from Pearson to weighted Pearson correlation entails that for binary weight-vectors (i.e. eq. 2), the formulation of equation 3 becomes identical to equation 2. This is because the weighted Pearson correlation uses the weighted mean and weighted variance in calculating the correlation. This effectively ignores time-points with a zero weight and fully include time-points with a weight set to one (for a longer discussion of the weighted Pearson correlation see (20)). Two sample illustrations of the concept of weight-vectors in the case of the sliding window method are given in Figures 1B-C.

The reformulation of *S*_*t*_ into *w*^*t*^ may at a first glance seem trivial. But the introduction of the weight-vector *w*^*t*^ allows us to generalize our formulation to many other DFC methods that are based on computing co-variance. As we will see, we can use *w*^*t*^ and estimate dynamic connectivity with the weighted Pearson correlation coefficient. This is because *w*^*t*^ does not have to be binary nor does it need its selection of supporting time points to be based on temporal proximity.

An extension of the sliding window approach is to use its tapered version. Here, *S*_*t*_ is not only a binary selection of time-points, but they are also weighted according to a statistical distribution (often a Gaussian distribution centered at *t*). The idea here is that time-points closer to *t* in time should have a greater impact on the estimate of the covariance at *t*. The tapered version of the sliding window method may easily be rewritten in the weight-vector formalism and we can use the weighted Pearson correlation to compute co-variance. This entails that all weight-values of *w*^*t*^ outside the tapered window are set to 0. Weight-values within the tapered window are set according to their probability given by the chosen distribution. Figures 1B-C show two examples of the tapered weight-vector centered at different points in time.

Instead of choosing *S*_*t*_ based on time-points which are temporally close, it is possible to choose *S*_*t*_ such that it emphasizes when the configuration of the state space (i.e. the multi-dimensional space spanned by all the ROIs included in the dFC analysis) is similar. This approach implies that time-points that are *spatially close* are selected to be included in *S*_*t*_. An example of this approach is given in (7) where we constructed binary weight-vectors based on spatial clustering (using k-means) of the state-space that was spanned by the BOLD fMRI time series from all brain ROIs. This spatial clustering approach can then be used to divide all time-points *t* = [1, …, *T*] into *K* different clusters. The assignment of clusters are then based on spatial dimensions across all ROIs across the brain. The individual weights in the weight-vector are set to 1 for all time-points belonging to the same cluster as time-point *t*, otherwise they are set to 0. Examples of binary weight vectors based on clustering of spatial patterns are shown in Figures 1B-C. The underlying assumption for dFC methods based on spatial clustering becomes clear when comparing the cluster assignments in Figures 1B-C with the signal amplitude of the ROIs shown in Figure 1A. Note that the weight-vectors at *t* = 10 and *t* = 14 partly overlap for both sliding window techniques, but their appearance are very different from the weight-vectors established by the spatial clustering approach. Importantly, the weight-vector formalism described here is able to capture the derivation of quite different DFC methods under the same umbrella and the results are easily visualized and illustrated as exemplified in Figure 1.

Similar to the tapered sliding window method, it is possible to extend the spatial clustering approach for DFC analysis so that continuous values for the weight-vector are employed when finding connectivity estimates that are based on spatial dimensions. In (20), we bypassed the clustering step by estimating the spatial distance between data time-points. The weight vector at *t* at index 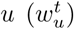 is computed by taking one over the Euclidean distance between the spatial dimensions at time-point *t* and time-point *u*. Thus, every data-point in time is assigned to a unique weight that can be described in the weight-vector formulation given above. An example of DFC based on spatial distance is given in Figures 1B-C and it can be seen that there are distinct similarities in weight values between the spatial clustering and spatial distance approach to DFC.

So far, we have illustrated how four different dynamic connectivity methods that all uses the Pearson correlation (2 binary, 2 continuous; 2 based on temporal proximity, 2 based on spatial proximity) can be formulated within a common framework. Each method can be formulated in terms of covariance estimates that are computed by weight-vectors and the weighted Pearson correlation technique.

Obviously, the choice of weight-vector has consequences for the estimate of dynamic connectivity. This is shown in Figures 2AB where the estimate of the correlation between ROI 1 and ROI 2 is visualized. This Figure shows how the resulting correlation is shaped by each of the different weight-vectors. The resulting difference in correlation across the four methods as well as which data-points that are included in the computation of co-variance is shown in Figure 2CD. When t=10 (Figure 2C) all four methods result in negative weighted Pearson correlation estimates, albeit with different magnitudes. At t=14 (Figure 2D), the four methods yields very different results. Both of the sliding window methods result in negative connectivity estimates whereas the spatial distance based weighted correlation is close to zero and the spatial clustering approach yields a positive correlation. In sum, the four methods give very different connectivity estimates but, importantly, the weight-vector formulation introduced here provide key insights to why this is so and it facilitates graphical comparisons between methods as illustrated in Figures 1 and 2.

Thus, we have shown that the weight-vector formulation can adequately capture all four DFC methods described so far. We can rephrase eq. 3 to include more than two spatial (ROIs) regions:

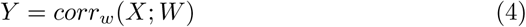

where *X* consists of *N* number of data channels, and *T* time points. *W* consists of *T* number of weight vectors. Finally, *Y* contains the resulting estimate of dynamic functional connectivity for each time-point and all channels.

**Figure 2.**
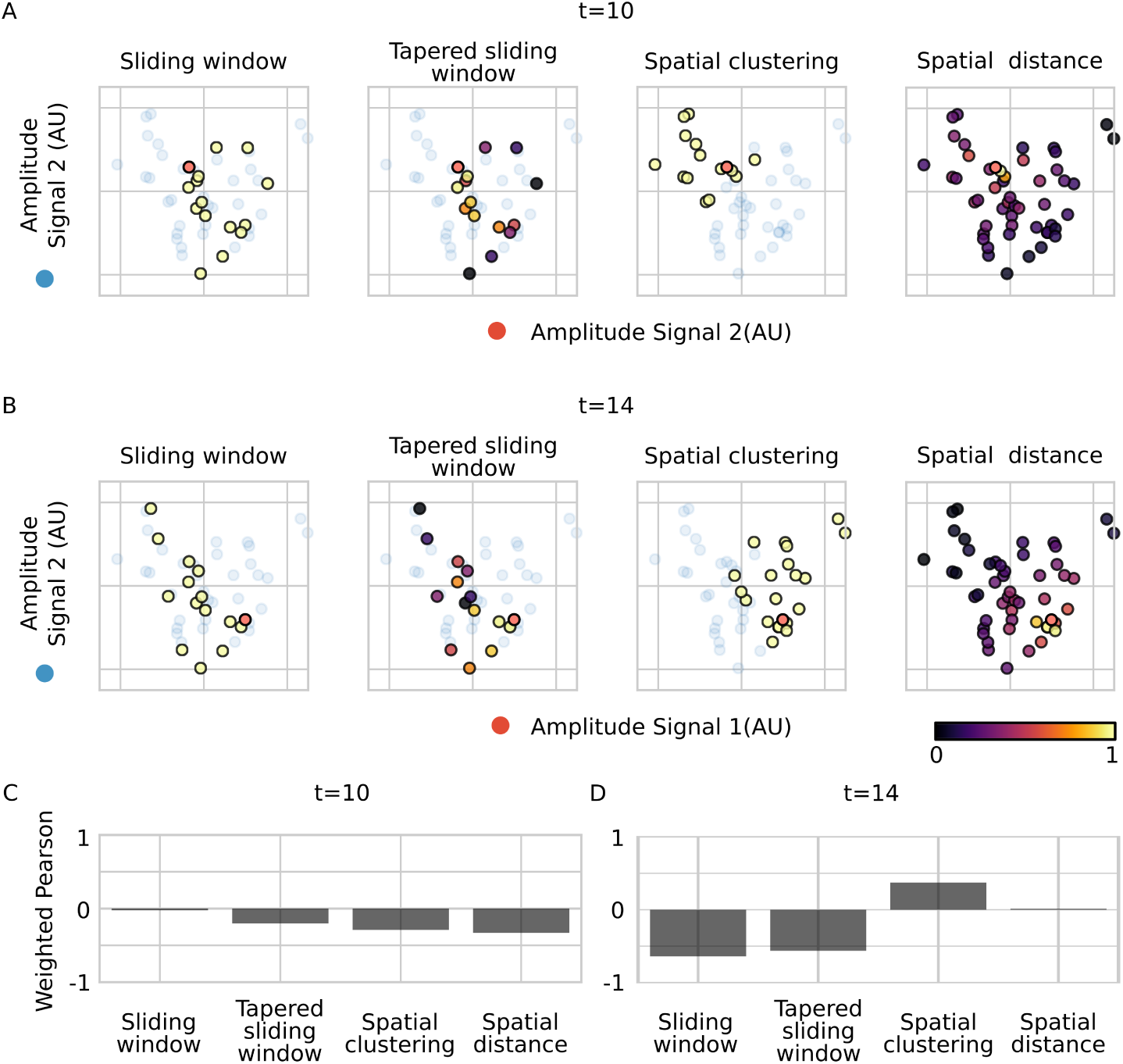
Visualization of BOLD fMRI signal values extracted from ROI 1 and ROI 2 shown in Figure 1A and their relationship to the estimated co-variance for the four DFC methods discussed in this paper. (A) Selection of data points that contribute to the weight-vectors for all four DFC methods at *t* = 10. Purple-yellow colours mark time-points which are given weights larger than 0. The red circles mark *t*. Faded blue circles mark data-points that were not included in the estimate of co-variance (i.e. their corresponding weights are 0). (B) Same as in (A) but at *t* = 14. (C) Weighted Pearson correlation between ROI 1 and ROI 2 using the four different methods. (D) same as in (C) but for *t* = 14.

The formulation given in eq. 4 does not, as it currently stands, generalize to all flavors of methods proposed to perform DFC. We will now show that eq. 4 can be extended, albeit more abstractly, to encompass other methods as well. In order to do that, we need to include a optional transformation of the raw data time-series and to allow for more flexibility in how different data time-series are related to each other. An example of how such an extension of eq. 4 can be formulated is to consider the DFC method proposed by (8). In Shine et al. 2015, the temporal derivative is used instead of the raw signal in their analysis (i.e. a transform is used). Further, the relationship between two brain areas was measured by multiplying the temporal derivatives of the signals with each other (i.e. a relational function), and then finally, a temporal averaging using a window function was carried out (a weight-vector). Thus, we can extend and generalize eq. 4 so that it can encapsulate the method proposed by the Shine et al. First, the temporal derivation of the raw data can be included by defining a transform function *U* that acts on the raw data *X* as:

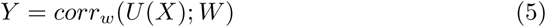

Obviously, when the raw signal is used, U is a simple function such that:

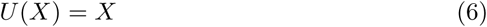

so nothing from the previous formulation is changed. In theory, *U* can be any mathematical transform applied to the raw data prior to the computation of co-variance. In the case of (8), *U* is the first temporal derivative of the raw data *X*:

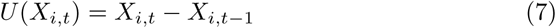

It is rather straightforward to accommodate other kinds of data transforms. For example, a transform that is commonly performed in the context of dynamic functional brain connectivity is a reduction of data dimensionality. In this case, *U* would be, for example, a principal component analysis transformation of the raw data.

The second modification of eq. 4 to accommodate the (8) method is to estimate the relationship between signal intensity time-courses. Instead of a covariance estimate, the temporal derivatives of the signal time-courses from two voxels or regions *i* and *j* are multiplied with each other and then divided by their standard deviations. Thus, we need to substitute *corr*_*w*_ in eq. 5 with a general relational function *R*:

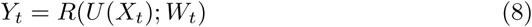

In the weighted Pearson correlation case, *R*(*U* (*X*); *w*) = *corr*_*w*_(*U* (*X*)). For the temporal derivative case described in (8), *R* becomes:

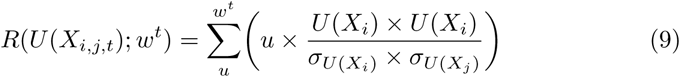

In the formulation above, we incorporate both the multiplication of the temporal derivatives and the moving average operation in *w*. We would like to point out that the temporal derivative method is just an example. *R* is a general relation function and it thus provide the means for any type of correlation or relation to be used. Possible types of relations that can be expressed in *R* are numerous.

For example, they may be non-parametric versions of the correlation coefficient (e.g. a weighted Spearman rank correlation), other regression models such as LASSO, Bayesian statistics to derive posterior distributions, different clustering models or machine learning techniques. Table 1 outlines how many currently used methods in the literature can be formulated with the formulation provided in eq 8.

**Table 1:** illustration how 9 different methods can be interpreted by specifying U R and W.

| Method | U | R | W |
| --- | --- | --- | --- |
| Sliding window | - | Weighted Pearson correlation | Window of length $w$. Binary weight, $w$ closest temporal points |
| Tapered Sliding window | - | Weighted Pearson correlation | Tapered window of length $w$. Continuous weight, $w$ closest temporal points |
| Spatial clustering | PCA (optional) | Weighted Pearson correlation | Clustering (e.g. K-means). Binary weight, closest spatial points |
| Spatial distance | - | Weighted Pearson correlation | Distance function (e.g. Euclidean). Continuous weight, closest spatial points |
| Jackknife correlation | - | Weighted Pearson correlation | All time points 1 to $T$ apart from $t$ |
| Temporal derivative | Derivative | Multiply and divide by multiplied standard derivations | Binary weight, $w$ closest temporal points |
| Hidden Markov model | - | Probability of hidden state | $t-1$ and $t$ |
| Wavelet coherence | Wavelet transform | Coherence | Window of length $w$. Binary weight, $w$ closest temporal points. |
| Dynamic conditional correlation | GARCH model | Exponentially weighted moving average | $\lambda$ parameter sets sensitivity previous time points. |

There are two additional considerations regarding the general formulation in equation 8. First, some methods for deriving estimates of DFC include a relational function *R* that necessitates certain choices of weight-vectors. For example, the hidden Markov model (HMM) is a possible choice of *R*. HMMs assume the Markov property which implies that only the data acquired at *t* − 1 is required in estimating the latent property at *t*. Thus, HMMs implicitly predetermine the shape of the weight-vectors, i.e. a binary weight vector where the only non-zero values are at *t* − 1 and *t*. However, despite the usage of methods that require predetermined choices of *W*_*t*_, researchers should be able to motivate why it is appropriate as the choice of weight-vectors is an underlying assumption.

The final consideration regarding the theoretical formulation given in equation 8 is related to the shape of *Y*_*t*_. It may be dependent on *U*, *W*_*t*_ and *R*. For example, the number of time-points will equal *T* in many methods but it may be shrunk for certain dFC methods (for example, the sliding window methods chops off (*w* − 1)/2 data-points at both the start and the end of the time-series). Three commonly used formats for the results of dynamic functional connectivity enclosed in *Y*_*t*_ are 3 dimensional (node x node x time, Figure 3A), 2 dimensional (component x time, Figure 3B), and 1 dimensional (time, Figure 3C). Note that the variable time is always present for all three outputs, since this is an essential property of dynamic functional connectivity analyses.

**Figure 3.**
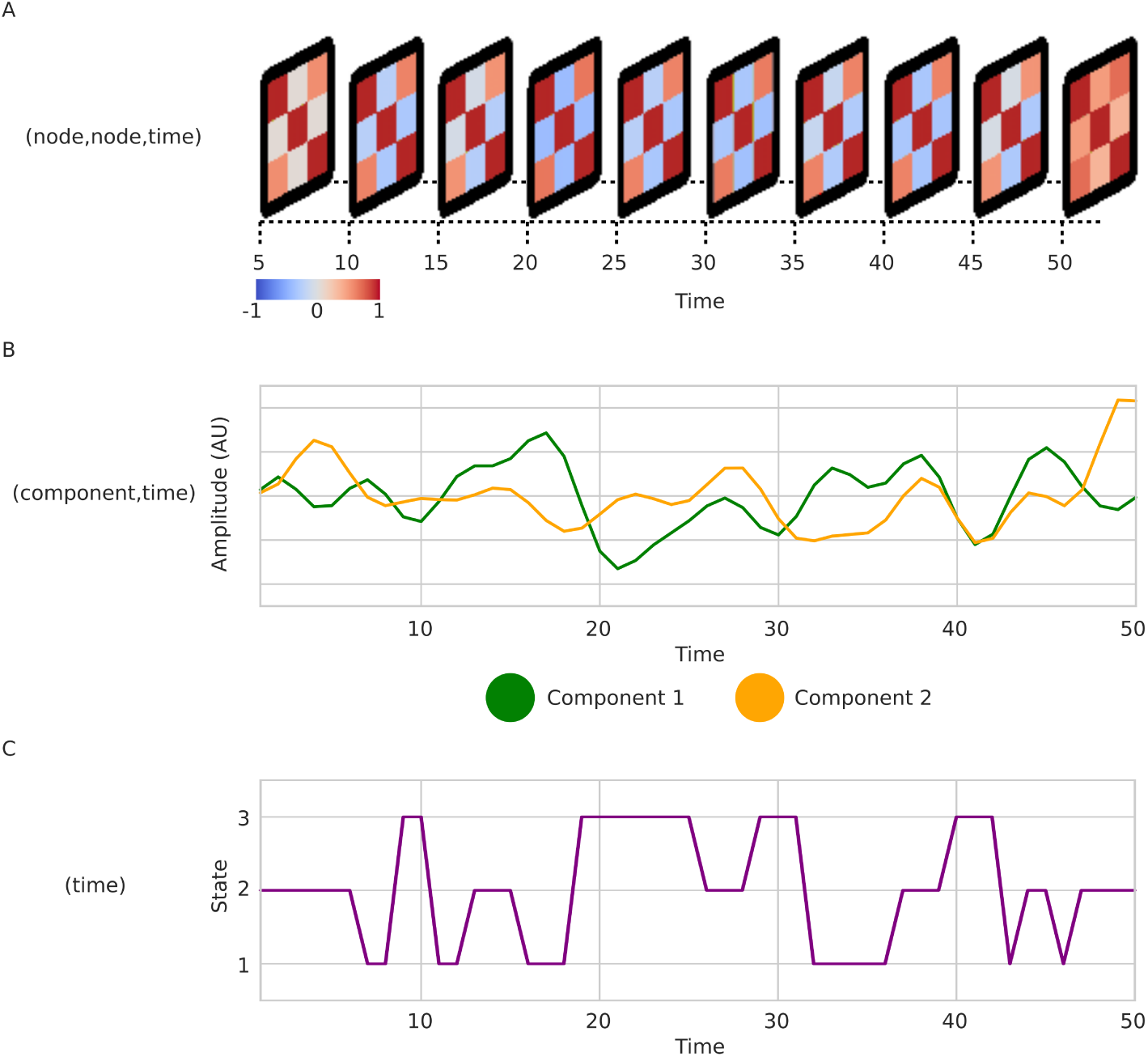
Different types of output from dynamic functional connectivity analysis using the data shown in Figure 1A. (A) A 3-dimensional representation (node,node,time). A graphlet stack plot showing connectivity matrices every 5 volumes. Correlations are derived using the spatial distance method (Euclidean distance). (B) A 2-dimensional representation (component,time) using 2 temporal ICA components. (C) A 1-dimensional representation (time). A vector of state assignments following k-mean clustering (*k* = 3).

## Discussion

In sum, we have shown that various methods to compute dynamic functional connectivity, which may seem vastly different on the surface, can nevertheless be unified into rather simple formulation. This provides a basis to understand and appreciate how we can compare the performance, results and underlying assumptions made for each method to compute DFC. To this end, we are proposing a general formulation given by *Y* = *R*(*U* (*X*; *W*)) in which most methods can be expressed.

The role of the formulation suggested here is to emphasize the assumptions different methods use and assisting in comparing methods. This is a critical aspect when evaluating the performance and caveats of different methods to compute DFC. The framework presented highlights three different assumptions that need to be made in all DFC methods: 1. The way in which the raw data is to be transformed before estimation of co-variance (*U*). 2. A relational function, *R* which quantifies the relationship between multiple data time-series. 3. Weighting vectors *W* which define time-points that are to be used to support the relational/covariance estimates. The first two assumptions are relatively trivial and are present in many data analysis problems. In the context of neuroimaging the choice of weight-vectors is perhaps more interesting as it is dictated by (1) the imaging modality (i.e. temporal and spatial resolution) and (2) how the brain works (i.e. speed of brain dynamics and the recurrence of spatial patterns).

While attempts to unify different but related problem formulations rarely contributes novel insight to the field, our hope is that our theoretical platform will assist researchers to understand the relationship between the many DFC methods proposed in the literature that in a sense all aim to tackle the same question. However, it can not be ruled out that some methods may fall outside the scope of the framework present here. The framework presented here is intended to assist method comparison and illustrate the underlying assumptions, not to be a rigid definition of dynamic functional connectivity that all methods must adhere to.

## Acknowledgements

This work was supported by the Swedish Research Council (grants no. 2016-03352 and 773 013-61X-08276-26-4) and the Swedish e-Science Research Center.

